# Neutralizing antibodies from early cases of SARS-CoV-2 infection offer cross-protection against the SARS-CoV-2 D614G variant

**DOI:** 10.1101/2020.10.08.332544

**Authors:** Cheryl Yi-Pin Lee, Siti Naqiah Amrun, Rhonda Sin-Ling Chee, Yun Shan Goh, Tze-Minn Mak, Sophie Octavia, Nicholas Kim-Wah Yeo, Zi Wei Chang, Matthew Zirui Tay, Anthony Torres-Ruesta, Guillaume Carissimo, Chek Meng Poh, Siew-Wai Fong, Wang Bei, Sandy Lee, Barnaby Edward Young, Seow-Yen Tan, Yee-Sin Leo, David C. Lye, Raymond T. P. Lin, Sebastien Maurer-Stroh, Bernett Lee, Wang Cheng-I, Laurent Renia, Lisa F.P. Ng

## Abstract

The emergence of a SARS-CoV-2 variant with a point mutation in the spike (S) protein, D614G, has taken precedence over the original Wuhan isolate by May 2020. With an increased infection and transmission rate, it is imperative to determine whether antibodies induced against the D614 isolate may cross-neutralize against the G614 variant. In this report, profiling of the anti-SARS-CoV-2 humoral immunity reveals similar neutralization profiles against both S protein variants, albeit waning neutralizing antibody capacity at the later phase of infection. These findings provide further insights towards the validity of current immune-based interventions.

**IMPORTANCE:** Random mutations in the viral genome is a naturally occurring event that may lead to enhanced viral fitness and immunological resistance, while heavily impacting the validity of licensed therapeutics. A single point mutation from aspartic acid (D) to glycine (G) at position 614 of the SARS-CoV-2 spike (S) protein, termed D614G, has garnered global attention due to the observed increase in transmissibility and infection rate. Given that a majority of the developing antibody-mediated therapies and serological assays are based on the S antigen of the original Wuhan reference sequence, it is crucial to determine if humoral immunity acquired from the original SARS-CoV-2 isolate is able to induce cross-detection and cross-protection against the novel prevailing D614G variant.

## OBSERVATION

Coronavirus disease 2019 (COVID-19) is the consequence of an infection by severe acute respiratory syndrome coronavirus 2 (SARS-CoV-2), which emerged in Wuhan, China, in December 2019 (1). The rapid expansion of the COVID-19 pandemic has affected 213 countries and territories, with a global count of more than 36 million laboratory-confirmed human infection cases to date (2). An inevitable impact of this pandemic is the accumulation of immunologically relevant mutations among the viral populations due to natural selection or random genetic drift, resulting in enhanced viral fitness and immunological resistance (3, 4). For instance, antigenic drift was previously reported in other common cold coronaviruses, OC43 and 229E, as well as in SARS-CoV (5–7).

In early March 2020, a non-synonymous mutation from aspartic acid (D) to glycine (G) at position 614 of SARS-CoV-2 spike (S) protein was identified (8). This variant, G614, rapidly became the dominant SARS-CoV-2 clade in Europe by May 2020, suggesting a higher transmission rate over the original isolate, D614 (8). *In vitro* and animal studies have also indicated that the G614 variant may have an increased infectivity, and may be associated with higher viral loads and more severe infections (8, 9). Notably, single point mutations have been shown to induce resistance to neutralizing antibodies in other coronaviruses, including SARS-CoV and Middle East respiratory syndrome (MERS-CoV) (10, 11). More importantly, mutations in the S protein of SARS-CoV-2 have been shown to induce conformational modifications that alter antigenicity (12, 13). Hence, determining any cross-neutralizing capability of antibodies developed against the earlier G614 variant is of paramount importance to validate the therapeutic efficacy of developing immune-based interventions.

Antibody profiling against the SARS-CoV-2 S protein was first assessed using plasma samples collected from COVID-19 patients (n=57) during the Singapore outbreak between January and April 2020, across the early recovery phase (median 31 days post illness onset [pio]) and a later post-recovery time point (median 98 days pio) (Table 1, Figure 1A and 1B). All patients showed a decrease in IgM response (Figure 1A), and a prolonged IgG response over time (Figure 1B). Although one recent study has demonstrated similar neutralization profiles against both D614 and G614 SARS-CoV-2 pseudoviruses, the virus clade by which the six individuals were infected with was not identified (9). According to Singapore’s SARS-CoV-2 clade pattern from December 2019 till July 2020, the D614G mutation only appeared in February 2020 (Figure 1C). Hence, with knowledge on the D614G status of a subset of COVID-19 patients (n=44 infected with D614, n=6 infected with G614, n=7 containing all other clades: O, S, L, V, G, GH or GR; Table 1, Figure 1C), the neutralizing capacity of these anti-SARS-CoV-2 antibodies was assessed using pseudotyped lentiviruses expressing the SARS-CoV-2 S protein tagged with a luciferase reporter as a surrogate of live virus (14). The neutralization EC50 values of each patient were interpolated from the respective dose-response neutralization titration curves (Table 2, Figure 1D and 1E, Supplementary figure 1). Notably, these antibodies were able to neutralize both SARS-CoV-2 D614 and G614 pseudoviruses at similar levels, despite having a significantly lower neutralization capacity at median 98 days pio in all COVID-19 patients (Figure 1D and 1E, Supplementary figures 1 and 2). Corroborating other studies, severe patients have a higher and persisting level of neutralizing antibodies as compared with both mild and moderate patients (Table 2, Supplementary figure 2) (15, 16). Of clinical importance, all the patients infected with either the D614 or G614 clade elicited a similar degree of neutralization against both D614 and G614 pseudoviruses (Figure 1F), suggesting that the D614G mutation does not impact the neutralization capacity of the elicited antibodies. Our results support the notion that the locus where the point mutation occurred is not critical for antibody-mediated immunity and may not have an impact on virus resistance towards antibody-based interventions (4, 17).

**Figure 1.**
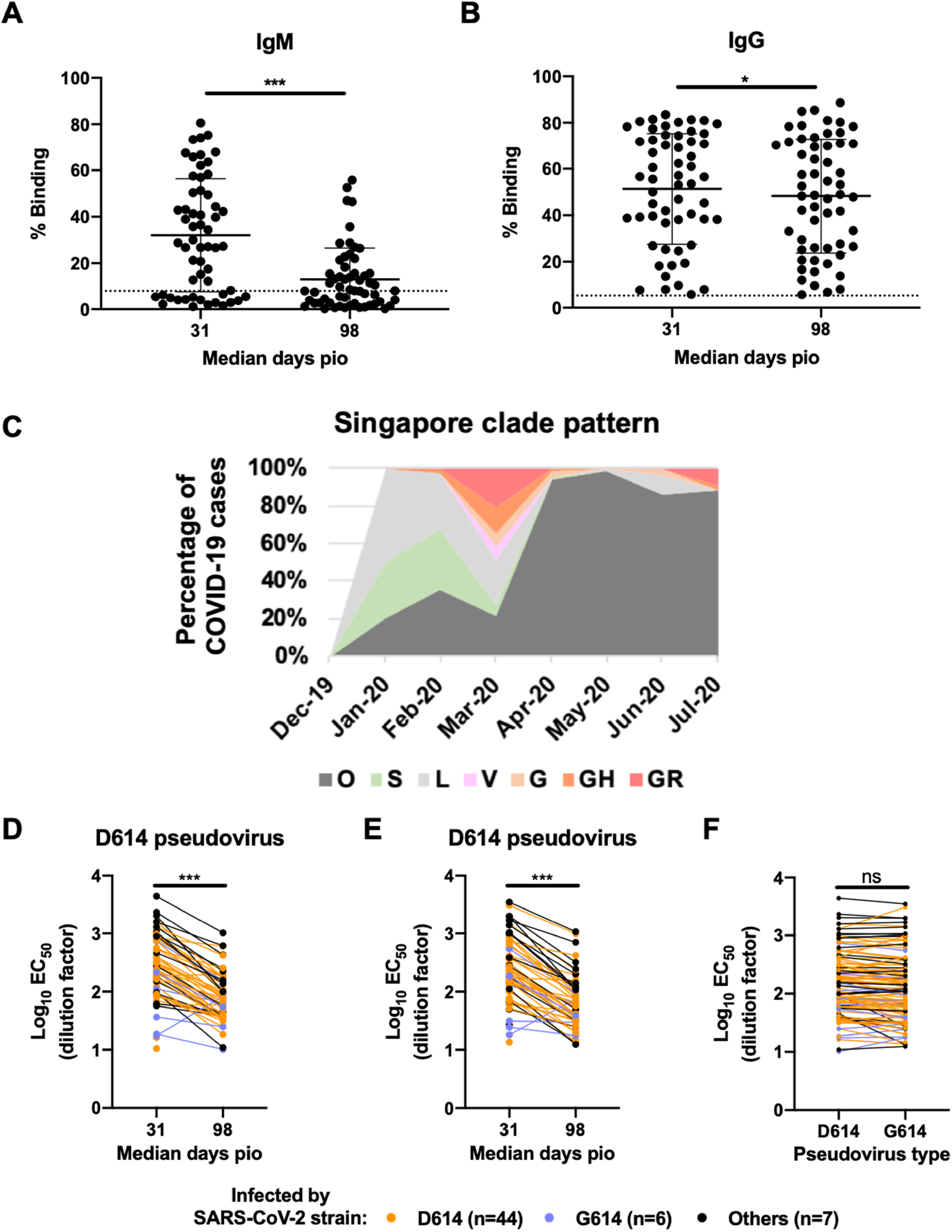
Timeline of events during the SARS-CoV-2 outbreak in Singapore, and the antibody profiles of COVID-19 patients and their neutralizing capacity against both D614 and G614 variants of SARS-CoV-2. Plasma samples of COVID-19 patients (n=57) at median 31 and median 98 days post-illness onset (pio) were assessed for anti-SARS-CoV-2 IgM and IgG antibody response. Plasma samples (1:100 dilution) were incubated with transduced HEK293T cells expressing SARS-CoV-2 spike protein, and (**A**) anti-IgM and (**B**) anti-IgG levels were quantified by flow cytometry. Data are shown as mean ± SD of two independent experiments. Dotted line indicates mean + 3SD of healthy controls (n=22). Statistical analysis was carried out with Wilcoxon signed-rank test (*P<0.05, ***P<0.001). (**C**) Percentage of COVID-19 cases during the Singapore outbreak from December 2019 to July 2020, segregated by the clade with which the patients were infected following GISAID clade nomenclature. **(D-F)** Anti-SARS-CoV-2 neutralizing antibodies were assessed using luciferase expressing lentiviruses pseudotyped with SARS-CoV-2 spike (S) protein of either the original strain, D614, or the mutant variant, G614. Log_10_ neutralization EC_50_ profiles against **(D)** D614 and **(E)** G614 pseudoviruses across both time points. Data represent the mean of two independent experiments and statistical analysis was carried out using paired *t* test (***P<0.001). (F) Comparison of log_10_ neutralization EC_50_ values between D614 and G614 pseudoviruses during both time points. Data represent the mean of two independent experiments and statistical analysis was carried out using paired ŕtest. All data point are non-significant (ns).

**Table 1.** Demographic and clinical information of COVID-19 patients.

|  |  |
| --- | --- |
| **Demographics** |  |
| Age, years | 45 (13) |
| Sex |  |
| Male | 38 (66.7%) |
| Female | 19 (33.3%) |
| Ethnicity |  |
| Chinese | 42 (73.7%) |
| Others | 15 (26.3%) |
| Comorbidities | 29 (50.9%) |
| Hyperlipidemia | 14 (24.6%) |
| Hypertension | 13 (22.8%) |
| Diabetes | 7 (12.3%) |
| Myocardial infection (history) | 5 (8.8%) |
| Others | 10 (17.5%) |
| **D614G infection status** |  |
| D614 | 44 (77.2%) |
| G614 | 6 (10.5%) |
| Others | 7 (12.3%) |
| **Clinical outcome (clinical severity; group)** |  |
| No pneumonia (0; mild) | 25 (43.9%) |
| Pneumonia, without hypoxia (1; moderate) | 19 (33.3%) |
| Pneumonia, with hypoxia (2; severe) | 13 (22.8%) |

Data represented as Mean (SD) or n (%). COVID-19: Coronavirus Disease 2019. Others: represent O, S, L, V, G, GH or GR clades.

**Table 2.**
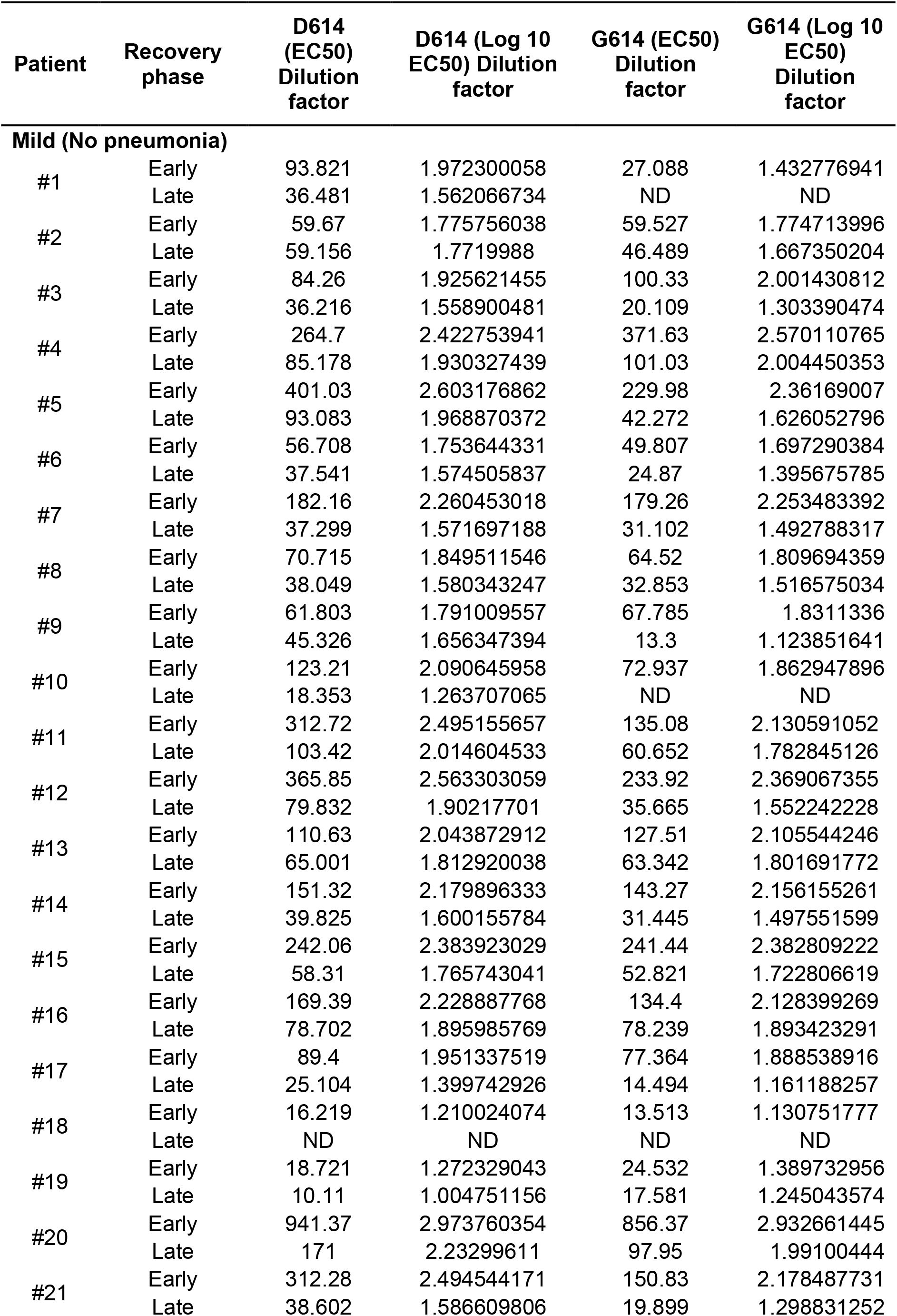
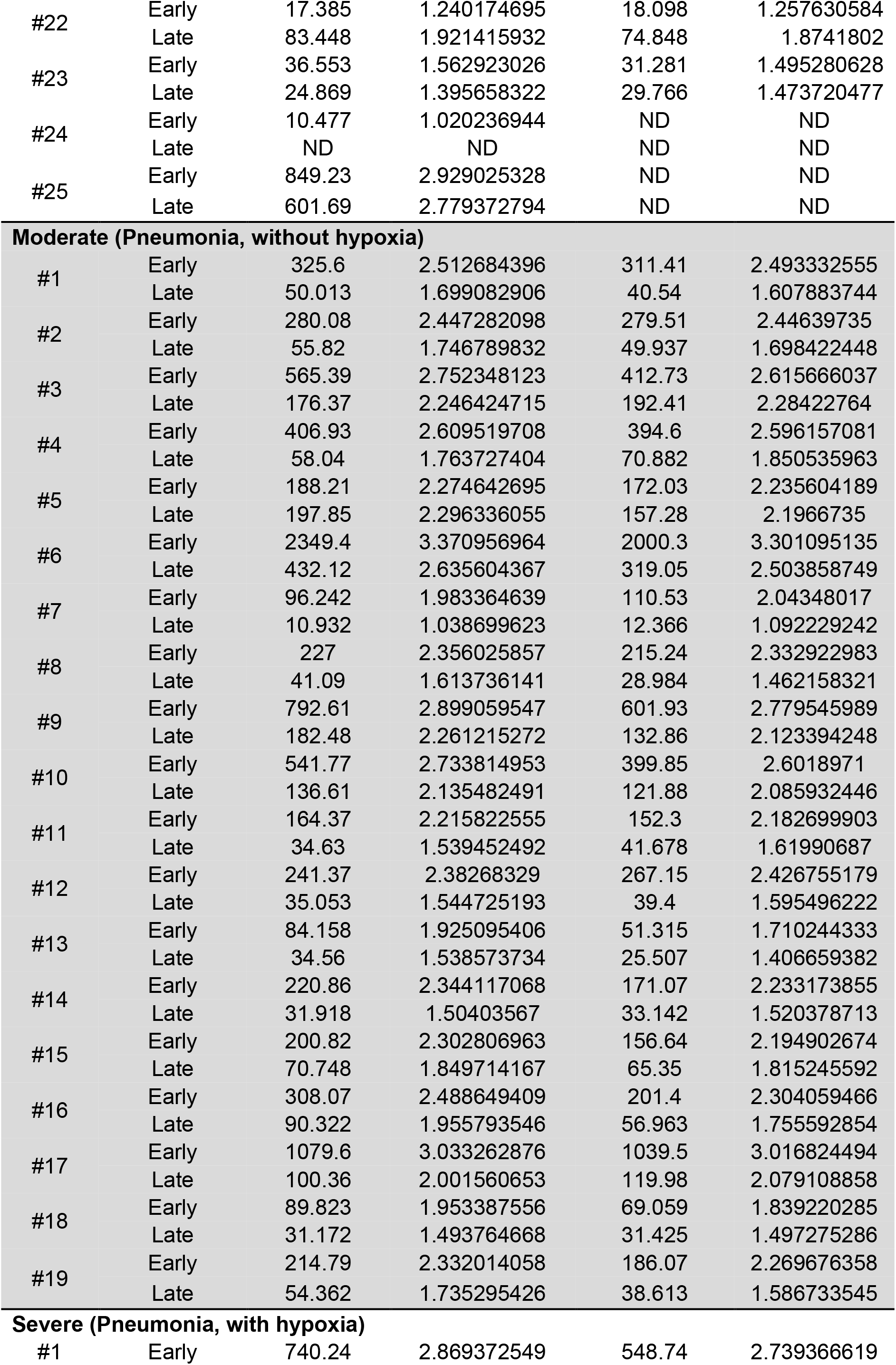
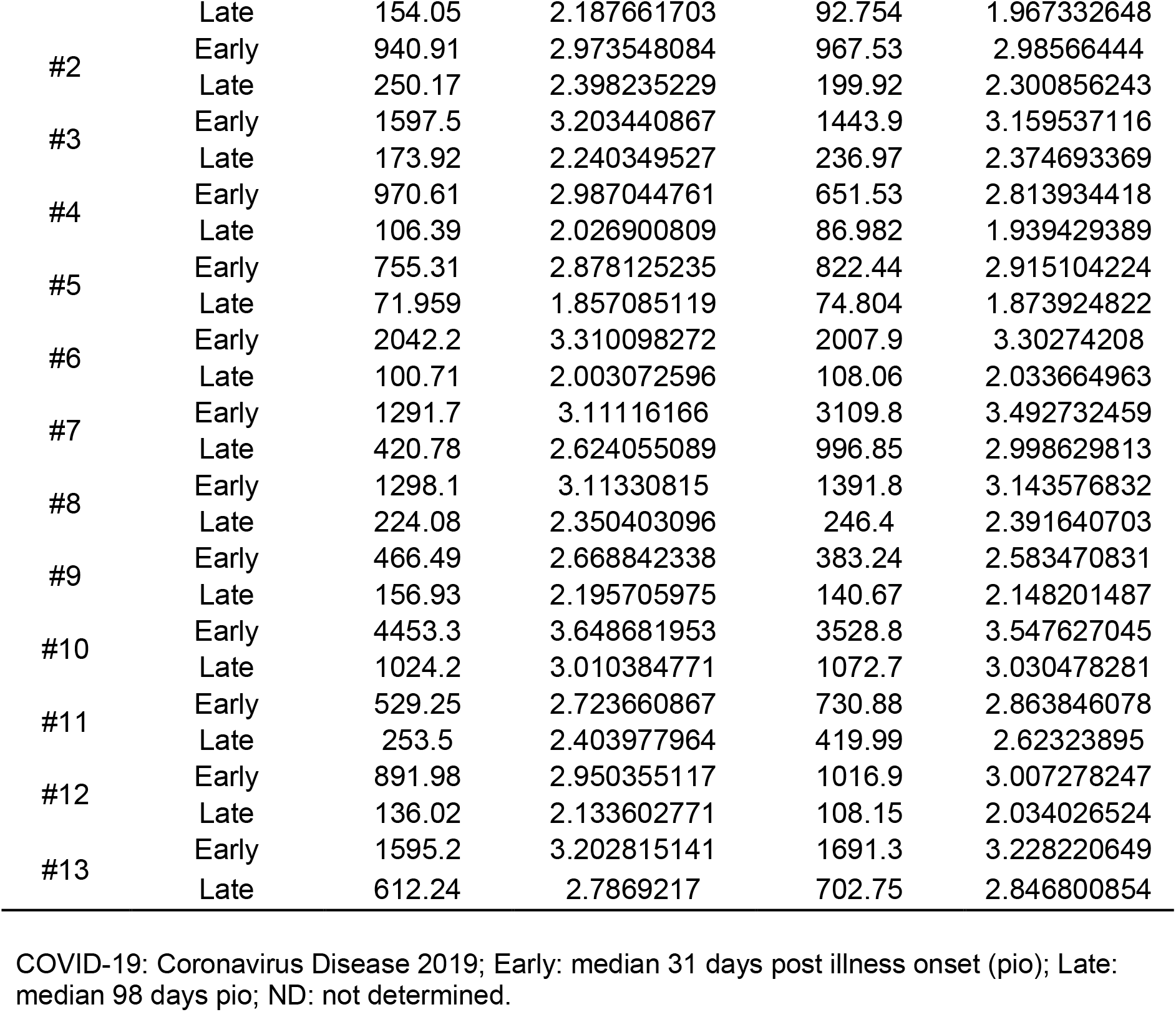
Neutralization EC50 values of COVID-19 patients.

The emergence of a new virus clade due to random mutations could heavily deter the therapeutic outcome of treatments and vaccines. Majority of the current immunoassays developed against SARS-CoV-2 are based on the S antigen of the original Wuhan reference sequence (18, 19). Moreover, pioneer batches of therapeutics and candidate vaccines were mostly designed based on earlier infections. As a result, mutations in the dominant variant sequence could potentially alter the viral phenotype and virulence, thereby rendering current immune-based therapies less efficient and effective (20, 21). Fortunately, a recent pre-print reported no observable difference in IgM, IgG and IgA profiles against either S variant in an antigen-based serological assay (22), providing preliminary findings on the effectiveness of current diagnostic approaches to detect SARS-CoV-2 G614 infections. In addition, determining the level of cross-reactivity is essential for immunosurveillance, as well as to identify broadly neutralizing antibodies or epitopes (23). Here, we confirm that cross-reactivity occurs at the functional level of the humoral response on both the S protein variants. Our results, together with the recent serological evaluation (22), strongly suggest that existing serological assays will be able to detect both D614 and G614 clades of SARS-CoV-2 with a similar sensitivity. However, it is of clinical relevance to assess if cross-reactivity between the variants may enhance viral infection when neutralizing antibodies are present at suboptimal concentrations (24). More importantly, further studies using monoclonal antibodies are necessary to validate the cross-reactivity profiles between both SARS-CoV-2 S variants.

Overall, our study shows that the D614G mutation on the S protein does not impact SARS-CoV-2 neutralization by the host antibody response, nor confer viral resistance against the humoral immunity. Hence, there should be negligible impact towards the efficacy of antibody-based therapies and vaccines that are currently being developed.

## Acknowledgements

The authors would like to thank the study participants who donated their blood samples to this project, and the healthcare workers caring for the COVID-19 patients. The authors also wish to thank Ding Ying and the Singapore Infectious Disease Clinical Research Network (SCRN) for their help in patient recruitment, and the staffs at the National Centre for Infectious Diseases (NCID) who assisted with data analysis on viral sequences and determination of the D614G status. The authors would also like to thank Professor Yee-Joo Tan (Department of Microbiology, NUS; Institute of Molecular and Cell Biology (IMCB), A*STAR) for kindly providing the CHO-ACE2 cells, and Dr. Brendon John Hanson (DSO National Laboratories, Singapore) for kindly providing the SARS-CoV-2 wildtype S protein.

## Author Contributions

CYPL, SNA, RSLC, YSG, TMM, and SO designed the experiments, acquired and analyzed the data, and wrote the manuscript. NKWY, ZWC, MZT, ATR and CMP acquired and analyzed the data, and wrote the manuscript. GC, SWF, WB, SL, SMS, BL and WCI analyzed the data and wrote the manuscript. BEY, SYT, YSL, DCL and RTPL designed and supervised sample collection. LR and LFPN conceptualized the study, analyzed the data and wrote the manuscript. All authors revised and approved the final version of the manuscript.

## References

1. Cohen J, Normile D. New SARS-like virus in China triggers alarm. Science. 2020;367(6475):234–5.

2. Worldometer. Dover, Delaware, U.S.A. 2020 [Available from: https://www.worldometers.info/coronavirus/?

3. Sevajol M, Subissi L, Decroly E, Canard B, Imbert I. Insights into RNA synthesis, capping, and proofreading mechanisms of SARS-coronavirus. Virus Research. 2014;194:90–9.

4. Grubaugh ND, Hanage WP, Rasmussen AL. Making Sense of Mutation: What D614G Means for the COVID-19 Pandemic Remains Unclear. Cell. 2020.

5. Ren L, Zhang Y, Li J, Xiao Y, Zhang J, Wang Y, et al. Genetic drift of human coronavirus OC43 spike gene during adaptive evolution. Scientific Reports. 2015;5(1):11451.

6. Chibo D, Birch C. Analysis of human coronavirus 229E spike and nucleoprotein genes demonstrates genetic drift between chronologically distinct strains. Journal of General Virology. 2006;87(5):1203–8.

7. Song H-D, Tu C-C, Zhang G-W, Wang S-Y, Zheng K, Lei L-C, et al. Cross-host evolution of severe acute respiratory syndrome coronavirus in palm civet and human. Proceedings of the National Academy of Sciences of the United States of America. 2005;102(7):2430.

8. Zhang L, Jackson CB, Mou H, Ojha A, Rangarajan ES, Izard T, et al. The D614G mutation in the SARS-CoV-2 spike protein reduces S1 shedding and increases infectivity. bioRxiv: the preprint server for biology. 2020:2020.06.12.148726.

9. Korber B, Fischer WM, Gnanakaran S, Yoon H, Theiler J, Abfalterer W, et al. Tracking Changes in SARS-CoV-2 Spike: Evidence that D614G Increases Infectivity of the COVID-19 Virus. Cell. 2020.

10. Sui J, Aird DR, Tamin A, Murakami A, Yan M, Yammanuru A, et al. Broadening of Neutralization Activity to Directly Block a Dominant Antibody-Driven SARS-Coronavirus Evolution Pathway. PLOS Pathogens. 2008;4(11):e1000197.

11. Tang X-C, Agnihothram SS, Jiao Y, Stanhope J, Graham RL, Peterson EC, et al. Identification of human neutralizing antibodies against MERS-CoV and their role in virus adaptive evolution. Proceedings of the National Academy of Sciences. 2014;111(19):E2018.

12. Eaaswarkhanth M, Al Madhoun A, Al-Mulla F. Could the D614G substitution in the SARS-CoV-2 spike (S) protein be associated with higher COVID-19 mortality? International Journal of Infectious Diseases. 2020;96:459–60.

13. Phan T. Genetic diversity and evolution of SARS-CoV-2. Infection, Genetics and Evolution. 2020;81:104260.

14. Poh CM, Carissimo G, Wang B, Amrun SN, Lee CY-P, Chee RS-L, et al. Two linear epitopes on the SARS-CoV-2 spike protein that elicit neutralising antibodies in COVID-19 patients. Nature Communications. 2020;11(1):2806.

15. Wang X, Guo X, Xin Q, Pan Y, Hu Y, Li J, et al. Neutralizing Antibody Responses to Severe Acute Respiratory Syndrome Coronavirus 2 in Coronavirus Disease 2019 Inpatients and Convalescent Patients. Clinical Infectious Diseases. 2020.

16. Zhao J, Yuan Q, Wang H, Liu W, Liao X, Su Y, et al. Antibody responses to SARS-CoV-2 in patients of novel coronavirus disease 2019. Clinical Infectious Diseases. 2020.

17. Barnes CO, West AP, Huey-Tubman KE, Hoffmann MAG, Sharaf NG, Hoffman PR, et al. Structures of Human Antibodies Bound to SARS-CoV-2 Spike Reveal Common Epitopes and Recurrent Features of Antibodies. Cell. 2020;182(4):828–42.e16.

18. Lee CY-P, Lin RTP, Renia L, Ng LFP. Serological Approaches for COVID-19: Epidemiologic Perspective on Surveillance and Control. Frontiers in immunology. 2020;11:879-.

19. Wang C, Gao Z, Shen K, Cao J, Shen Z, Jiang K, et al. Safety and efficiency of endoscopic resection versus laparoscopic resection in gastric gastrointestinal stromal tumours: A systematic review and meta-analysis. European Journal of Surgical Oncology. 2020;46(4, Part A):667–74.

20. Sanjuán R, Nebot MR, Chirico N, Mansky LM, Belshaw R. Viral Mutation Rates. Journal of Virology. 2010;84(19):9733.

21. Ojosnegros S, Beerenwinkel N. Models of RNA virus evolution and their roles in vaccine design. Immunome research. 2010;6 Suppl 2(Suppl 2):S5–S.

22. Klumpp-Thomas C, Kalish H, Hicks J, Mehalko J, Drew M, Memoli MJ, et al. D614G Spike Variant Does Not Alter IgG, IgM, or IgA Spike Seroassay Performance. medRxiv. 2020:2020.07.08.20147371.

23. Hicks J, Klumpp-Thomas C, Kalish H, Shunmugavel A, Mehalko J, Denson J-P, et al. Serologic cross-reactivity of SARS-CoV-2 with endemic and seasonal Betacoronaviruses. medRxiv: the preprint server for health sciences. 2020:2020.06.22.20137695.

24. Arvin AM, Fink K, Schmid MA, Cathcart A, Spreafico R, Havenar-Daughton C, et al. A perspective on potential antibody-dependent enhancement of SARS-CoV-2. Nature. 2020;584(7821):353–63.

